# *Parvimonas micra*, an oral pathobiont associated with colorectal cancer, epigenetically reprograms human primary intestinal epithelial cells

**DOI:** 10.1101/2022.05.14.491935

**Authors:** Emma Bergsten, Denis Mestivier, Francoise Donnadieu, Thierry Pedron, Caroline Barau, Landry Tsoumtsa Meda, Amel Mettouchi, Emmanuel Lemichez, Olivier Gorgette, Mathias Chamaillard, Amaury Vaysse, Stevenn Volant, Abiba Doukani, Philippe J. Sansonetti, Iradj Sobhani, Giulia Nigro

## Abstract

Recently, an intestinal dysbiotic microbiota with enrichment in oral cavity bacteria has been described in colorectal cancer (CRC) patients. Here we characterized and investigated one of these oral pathobionts, the Gram-positive anaerobic coccus *Parvimonas micra.* We identified two phylotypes (A and B) exhibiting different phenotypes and adhesion capabilities. We observed a strong association of phylotype A with CRC, with its higher abundance in feces and in tumoral tissue compared with the normal homologous colonic mucosa, which was associated with a distinct methylation status of patients. By developing an *in vitro* hypoxic co-culture system of human primary colonic cells with anaerobic bacteria, we showed that *P. micra* phylotype A alters the DNA methylation profile promoters of key tumor-suppressor genes, oncogenes, and genes involved in epithelial-mesenchymal transition. In colonic mucosa of CRC patients carrying *P. micra* phylotype A, we found similar DNA methylations alterations, together with significant enrichment of differentially expressed genes in pathways involved in inflammation, cell adhesion, and regulation of actin cytoskeleton, providing evidence of *P. micra* possible role in the carcinogenic process.

## Introduction

Colorectal cancer (CRC) is a multifactorial disease due to various genetic and environmental factors that contribute to tumor formation and disease development. Genetic predispositions for CRC, such as Lynch syndrome (also called human no polyposis colonic cancer-HNPCC) or familial adenomatous polyposis (FAP), only accounts for a minority of CRC, representing less than 5% of all cases^1^. The overwhelming majority of CRCs are sporadic cancers, i.e., the cause is unknown. Several lifestyle environmental factors increase the risk of CRC including physical inactivity^2^, overweight^3^ and high consumption of red meat, processed meat and unsaturated fatty acids^4^. The identification of lifestyle factors supports the hypothesis that the increase in CRC incidence is strictly related to environmental changes. Among environmental factors, the role of microorganisms in cancer has been increasingly recognized and a global imbalance, called dysbiosis, of gut microbiota compared to healthy individuals was observed in several cancers such as biliary, hepatic, breast cancers, or CRC^5^. Thus, the gut microbiota has emerged as an important carcinogenic environmental factor, in particular for CRC, due to the gut microbiota proximity and constant crosstalk with the intestinal epithelium^6^. Different bacterial species, as *Bacteroides fragilis*, *Fusobacterium nucleatum (F. nucleatum)*, pathogenic strains of *Escherichia coli* (CoPEC), or *Streptococcus gallolyticus*, have been associated with CRC development and mechanistic studies have tried to understand roles of these bacteria in carcinogenesis^7–10^. Recently, our team and others have identified, by metagenomic studies from humans’ feces, not only the intestinal bacterium *F. nucleatum* associated with CRC but also different bacteria belonging to the oral microbiota, such as *Gemella morbillorum*, *Solobacterium moorei*, *Porphyromonas gingivalis, and Parvimonas micra* (*P. micra*)^11–13^. The overabundance of these bacteria in the colonic niche may be an etiological factor in CRC^14^, but their role in carcinogenesis remain to be established. Here, we chose to focus on the candidate bacterium, *P. micra*.

*P. micra*, the only described species within its genus, is a Gram-positive anaerobic coccus and commensal of the oral cavity^15^. *P. micra* could be considered as a pathobiont^16^ because it is often isolated from oral polymicrobial infections associated with periodontitis^17^, from superficial polymicrobial infections, including wounds, ulcers, and skin abscesses^18–22^, as well as from deep tissue infections in the brain, lung and reproductive organs^23^. *P. micra* has been poorly studied due to difficulties in its cultivation and laboratory identification by traditional methods and the polymicrobial nature of the infective zones from which it is usually isolated.

The aim of this work was to better characterize *P. micra*, define its carriage in CRC patients, analyze *in vitro* the nature of the crosstalk between this oral pathobiont and colonic epithelial cells, and determine if the observed impact on primary epithelial cells were relevant in CRC patients.

## Results

### *P. micra* exists in two phylotypes with phenotypic and genetic diversity

To explore *P. micra* diversity, we began to study the twenty-seven clinical isolates already available in biobanks, which were collected from various types of infections (blood, systemic, and oral cavity) (Table S1). *P. micra* colonies on blood agar plates appeared heterogeneous. A contact hemolytic activity on blood agar plates was observed in twenty isolates (20/27) and *P. micra* colonies were found to be either “compact” (7/27) or “non-compact” (20/27) (Figure 1A, Figure S1A). In addition, sedimentation assays on two representative isolates of each phenotype: ATCC 33270 (“non-compact”) and HHM BlNa17 (“compact”), (named hereafter *PmA* and *PmB*), showed differences in the aggregation rate (Figure S1B). Phenotypical differences observed among the twenty-seven isolates could reflect genetic diversity. Therefore, we performed the full-length sequencing of the 16S rDNA on these isolates and compared them with the twenty-two rDNA16S sequences available on NCBI database. This revealed two distinct phylogenetic groups: A and B (Figure 1A). *Finegoldia magna (F. magna)*, the most closely related bacterial species to *P. micra*, was used to set the root of the phylogenetic tree. To confirm the existence of the two phylotypes, we have clustered whole genome sequences, available on NCBI, of ten of the 22 *P. micra* strains used for the 16S rDNA alignment, on the basis of the presence/absence of genes. This analysis confirmed the 16S rDNA clustering and the existence of phylotypes A and B (Figure S1C). Interestingly, all *P. micra* isolates from phylotype A were hemolytic whereas isolates belonging to phylotype B were not. Likewise, most phylotype B isolates were “compact” whereas phylotype A isolates were not. These data provide genetic evidence of two phenotypically distinct phylotypes in *P. micra* whose respective impact on colonocytes and distribution in CRC had to be defined.

**Figure 1:**
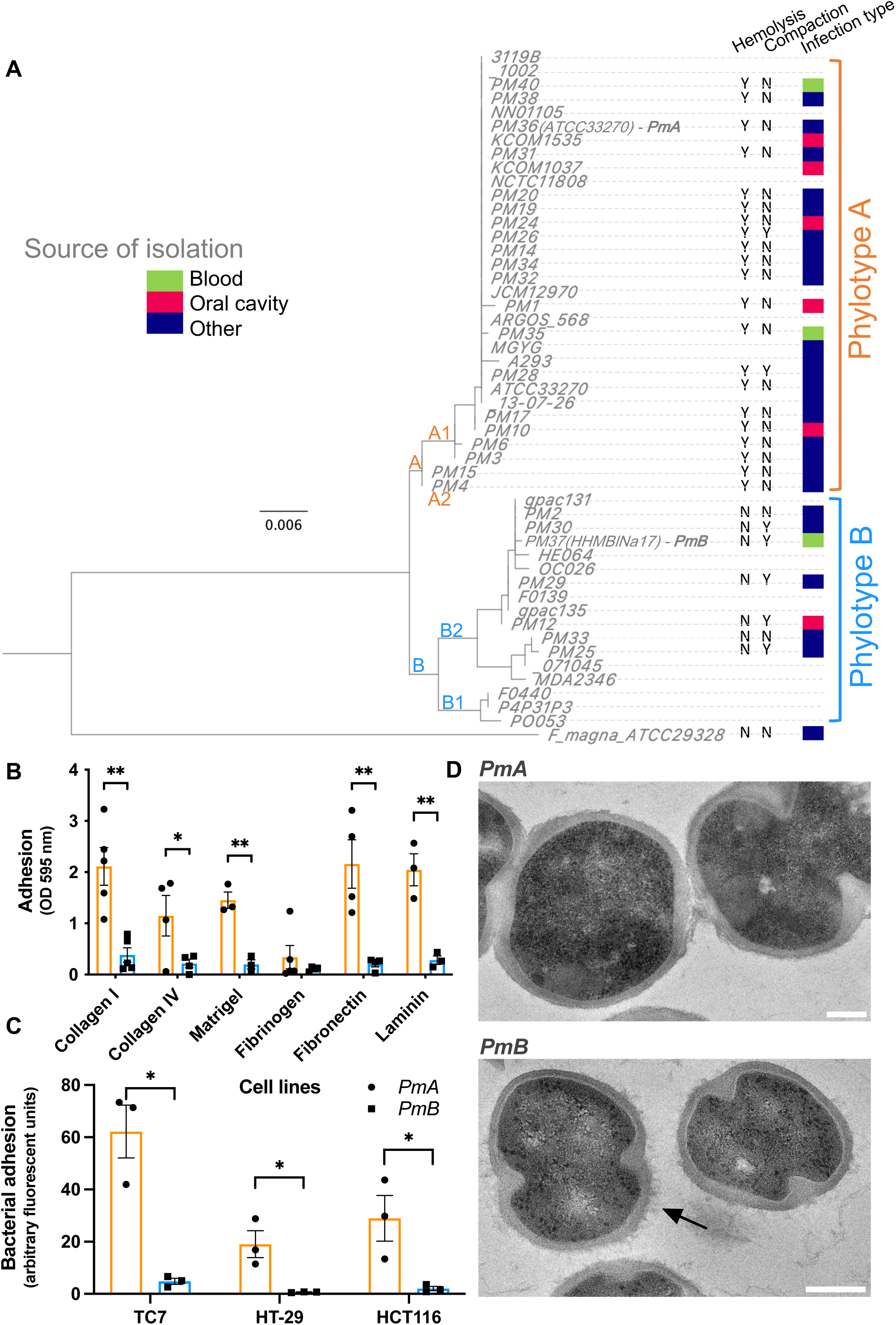
*P. micra* exists in two phylotypes with phenotypic and genetic diversity. A) Phylogenetic tree based on full length 16S rDNA sequences of 27 *P. micra* clinical isolates (named *PMX)* and 22 reference sequences from the NCBI database. Root: *Finegoldia magna* ATCC 29328. The infection type from which the 27 clinical isolates are derived are indicated by a color code; haemolysis and colony compaction abilities are indicated as present (Y) or absent (N). B) Adhesion capacity of *P. micra* ATCC 33270 (*PmA)* and HHM BlNa17 (*PmB)* to extracellular matrix proteins. Optical density at 595 nm represents measurement of bacterial adhesion. C) Adhesion capacity of *PmA* and *PmB* to different human colonic cell lines (TC7, HT-29, and HCT116) after 1 hour of co-culture. Bacterial adhesion was quantified by the analysis of fluorescent images of cells co-cultured with the two phylotypes and is reported as arbitrary fluorescent units (x100). Data are represented as mean ± SEM from three independent experiments. Mann-Whitney test **p<0.01; *p<0.05. D). Transmission electron microscopy on ultrafine sections of *PmA* (top) and *PmB* (bottom). Spiky surface structures (arrow) were observed only in *PmB* and not in *PmA*. Scale: 200 nm. See also Figure S1 and Table S1.

Clinical studies using 16S rDNA sequencing from tissue had shown the presence of *Parvimonas* genus sequences in tumoral and normal homologous colonic mucosa of CRC patients, suggesting that this microorganism may be able to adhere to colonocytes or to the colonic cellular matrix ^24, 25^. We therefore measured the ability of a representative isolate from each phylotype, *PmA* (phylotype A) and *PmB* (phylotype B), to adhere to different extracellular matrix (ECM) proteins and three human colonic cell lines*. PmA* adhered to the ECM proteins collagen I, collagen IV, fibronectin and laminin, as well as to Matrigel® but not to fibrinogen, whereas *PmB* did not adhere to any of these ECM proteins (Figure 1B). Similarly*, PmA* adhered 6-, 2-, and 3-fold more to TC7, HT-29, and HCT116, respectively as compared to *PmB* (p<0.05) (Figure 1C), indicating heterogeneity of adhesion capacity in *P. micra* phylotypes. To further investigate the adhesion capacity of phylotypes, we focused on *P. micra* isolated from the oral cavity, its endogenous niche. The isolate *Pm12*, belonging to phylotype B, neither adhered to Matrigel® (Figure S1D) nor to the human colonic cell line TC7 (Figure S1E), while those of phylotype A, *Pm1*, *Pm24*, and *Pm10,* did (p<0.01).

To explore whether the aggregative phenotype could be due to specific bacterial surface structural differences, ultrathin sections of the two phylotypes were analyzed by transmission electron microscopy (TEM). *PmB* (non hemolytic, “compact”) presented spiky surface structures rarely observed in *PmA* (hemolytic, “non-compact”) (Figure 1D). Similar spiky structures were observed in phylotype B oral isolate *Pm12,* but not in the phylotype A oral isolate *Pm10* (Figure S1F).

These data show that two phenotypically and genetically different phylotypes can be described for *P. micra*.

### The adherent *P. micra* phylotype A is associated with CRC

Differences in the *in vitro* adhesion capacity of the two phylotypes might reflect a different colonization ability *in vivo*. To assess this hypothesis, we first improved the resolution of our previous whole genome sequencing (WGS) metagenomic analysis^26^, focusing on the taxonomic assignation of raw data, in order to precisely investigate the occurrence and abundance of *P. micra* of CRC patients’ feces. We observed an enrichment of *P. micra* occurrence in the feces of CRC patients as compared to control healthy individuals (27% versus 1.1%). Three taxa of *P. micra* were detected: *’Parvimonas micra*’ from phylotype A, ’*Parvimonas* sp. oral taxon 393 - F0440’ and ’*Parvimonas* sp. oral taxon 110 - F0139’ from phylotype B. All *Parvimonas* positive patients carried the phylotype A in their feces, except in one patient where reads of both phylotypes were detected.

We then analyzed *Parvimonas* carriage in feces of CRC patients and control individuals, as well as from paired tumor and normal homologous tissues (adjacent to the tumor), using 16S rDNA sequencing of the V3-V4 region. As expected, *Parvimonas* prevalence in feces was enriched in CRC patients (81%), with no differences within the stages (international TNM (tumor-node-metastasis) staging system), compared with control individuals (60%) (Figure 2A). In 78% of CRC patients, *Parvimonas* sequences were detected in both tumoral and normal homologous samples with a significant abundance enrichment in the tumoral tissues (2.20% versus 0.53% of bacterial reads, respectively; p<0.01) (Figure 2B). Prevalence of *Parvimonas* in tissues showed no difference according to the TNM staging, with 74% in stages I-II versus 84% in stages III-IV and a respective abundance of 1.34% and 1.41% of bacterial reads (Figure 2B).

**Figure 2:**
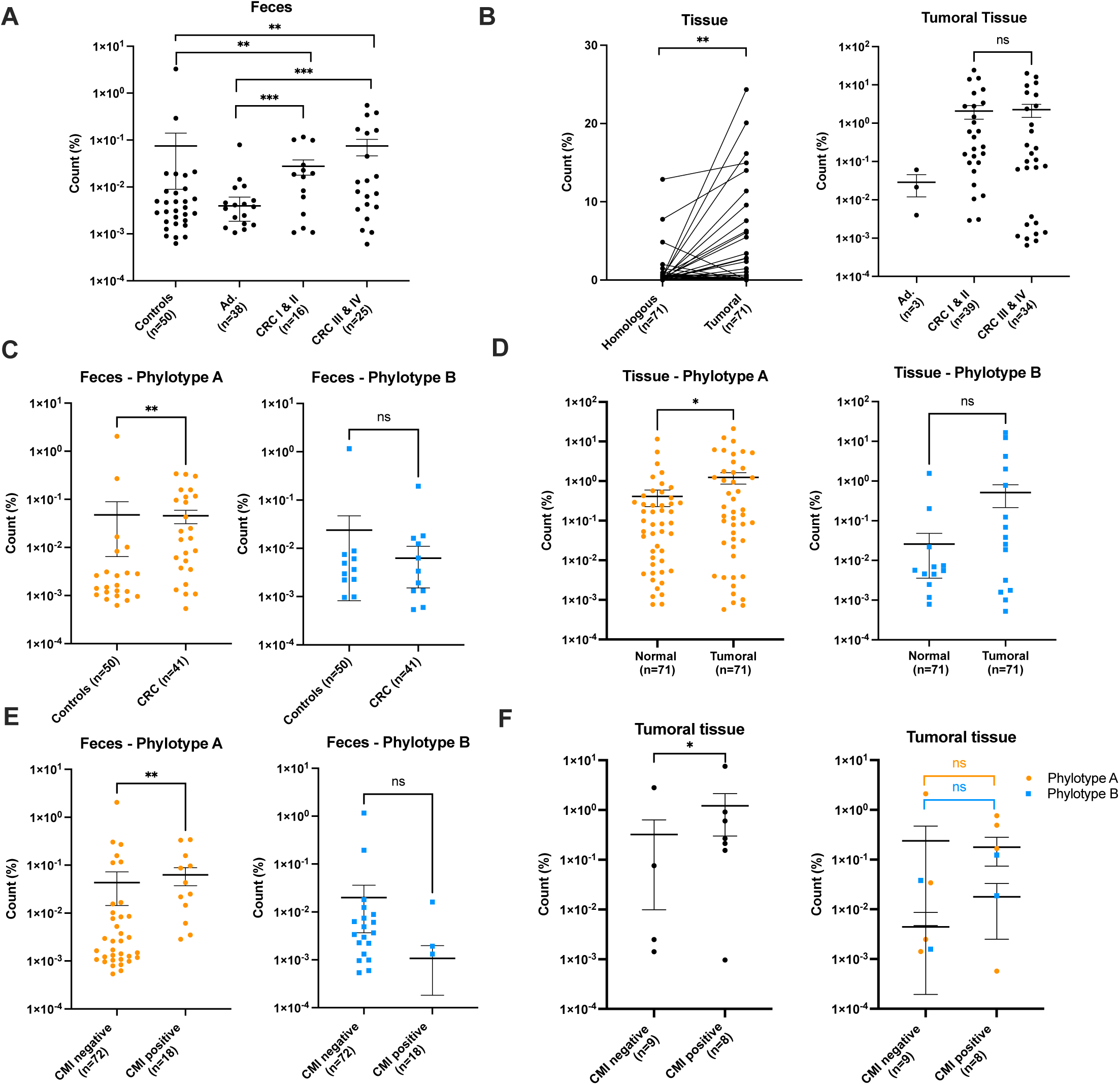
*P. micra* phylotype A is associated with CRC. *Parvimonas* abundance, determined by 16S rDNA sequencing of the V3-V4 region, in controls, adenoma (Ad.), or sporadic CRC patients at early (CRC I & II) and late (CRC III & IV) stages of carcinogenesis in A) feces (Test Mann-Whitney ***p<0.001; **p<0.01) or B) mucosa: on the left, normal homologous mucosa versus tumoral tissue of CRC patients (Wilcoxon’s Signed Wilcoxon Rank Test for paired samples **p<0.01); on the right, the same results only in tumoral tissue and according to CRC TNM classification. C) *P. micra* A and B phylotypes abundances in control and CRC patients’ feces, or D) associated with normal homologous mucosa and tumoral tissue of CRC patients. *P. micra* phylotypes association with CMI methylation score of the WIF1, PENK, and NPY genes promoters in E) controls and sporadic CRC patients’ feces, or F) associated with tumoral tissue of CRC patients. “n*”* represents the total number of analyzed samples and results from *Parvimonas* positive samples are expressed as a percentage of count (number of sequences assigned to *Parvimonas* per number of total bacterial sequences X100) mean +/- SEM. Statistical analysis of C, D, E, and F using the Mann-Whitney test **p<0.01, *p<0.05. See Table S2.

The V3-V4 regions of the 16S rDNA was then analyzed to discriminate between *Parvimonas* phylotypes A and B. Forty-four percent of control individuals carried the phylotype A in their feces as compared to 61% of the CRC patients (p<0.01), whereas no difference in the prevalence of the phylotype B was observed (22% and 27%, respectively) (Figure 2C). An increase in the abundance of the phylotype A, but not of the phylotype B, was also observed in tumor tissues compared to homologous normal tissues (p<0.05) (Figure 2D).

Previously, we demonstrated a strong correlation between *Parvimonas micra* carriage in feces and a positive cumulative methylation index (CMI) in the blood based on the methylation levels of promoters in the WIF1 gene involved in carcinogenesis and PENK and NPY, two neuromediators genes^13^. Thus, we wondered whether the phylotypes herein described, were correlated with the CMI. Analysis of 16S rDNA sequencing data from feces showed that phylotype A was enriched in CMI positive patients (p<0.01) but this was not the case for phylotype B (Figure 2E). *Parvimonas* carriage in tissues was also associated with a positive CMI (p<0.05), although the number of samples was not sufficient to discriminate between the two phylotypes (Figure 2F).

These results indicate that *Parvimonas micra* phylotype A sub-species is associated with CRC and a positive CMI.

### *P. micra*’s impact on human colonic primary epithelial cells

Colorectal tumor is defined as a group of anarchically proliferating cells with DNA mutations and aberrant DNA methylation. To determine whether *P. micra* has an impact on host cells physiology, we first developed a compatible co-culture model between this oxygen-sensitive anaerobic microorganism and primary human colonic epithelial cells. Colonic samples were obtained from normal site (distant from tumor tissue) of two colonic tumor resections. Colonic crypts were isolated and cultured as organoids (Figure 3A), allowing amplification of initial material through stem cell proliferation. To develop a system where bacteria could be in contact with the apical pole of primary cells, reflecting a physiological situation, organoid fragments were seeded onto a Transwell® insert system and grown to confluency up to a polarized monolayer (Figure 3B), as previously described^27^. Cell growth and formation of a tight monolayer were monitored over time by phase-contrast microscopy and transepithelial electrical resistance (TEER) measurements (Figures 3C and D). Under these culture conditions, monolayers were fully confluent three days after organoids fragments were seeded on the inserts (Figure 3D).

**Figure 3:**
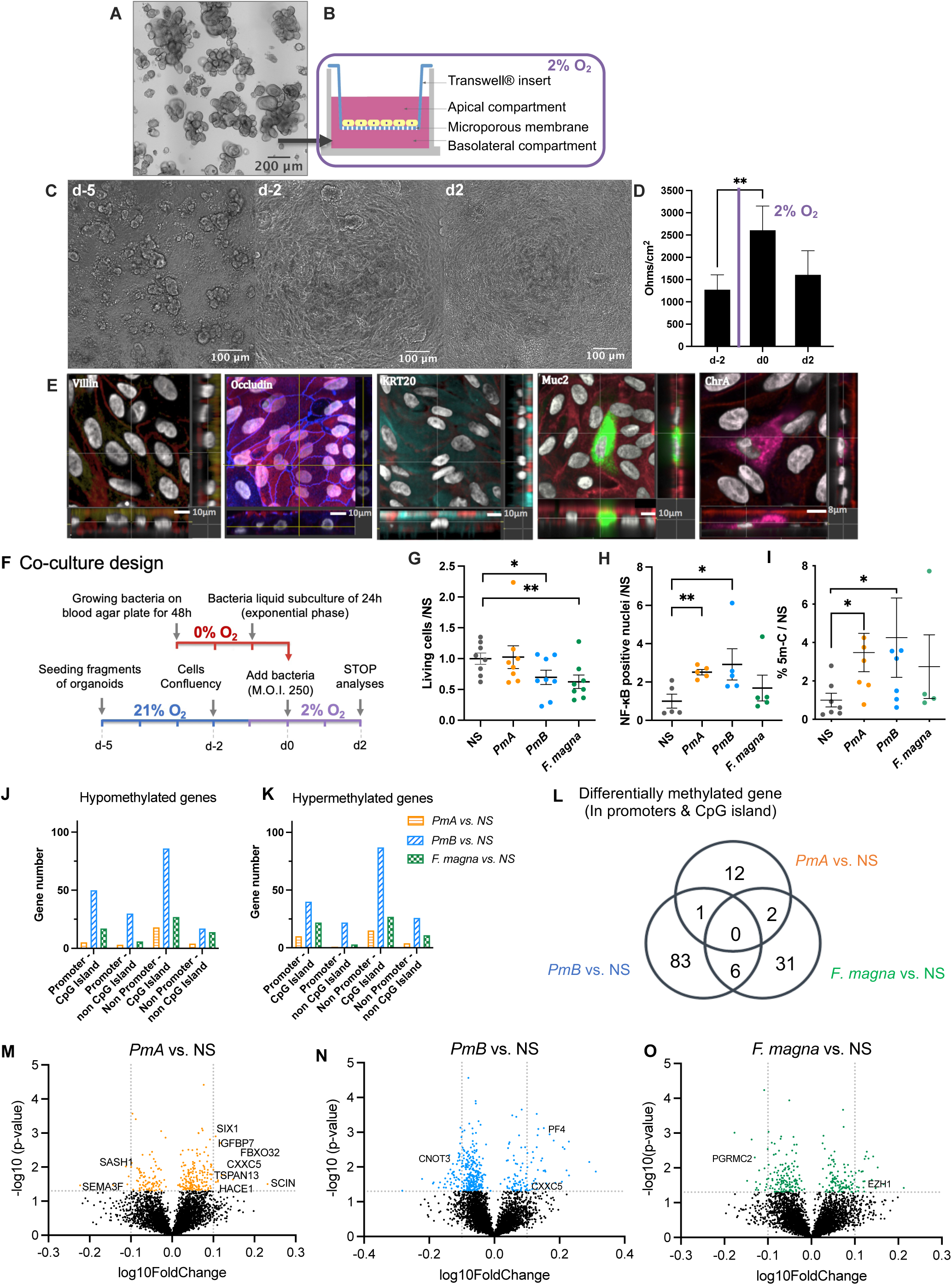
*P. micra* impact on human colonic primary epithelial cells. A) Colonic organoids obtained from normal tissue of CRC patients and cultured in Matrigel® with growth factors-enriched medium. B) Schematic representation of Transwell® permeable insert used to cultivate monolayers of primary colonic epithelial cells derived from organoids. C) Representative phase contrast images of organoid monolayers at seeding (d-5), after 3 days of culture in aerobic conditions (d-2) and after 7 days of culture, including 4 days in aerobic conditions and 3 days in hypoxic conditions (d2). D) Transepithelial electrical resistance (TEER) at different days of culture. TEER experiments were performed on cells from two donors, for at least 4 experiments per donor. Results are expressed as a mean of Ohm per cm^2^ +/- SEM. Mann-Whitney test **p<0.01. E) The differentiation of the monolayers after 4 days in aerobic conditions and 3 days in hypoxic conditions (2% O_2_) was assessed by confocal microscopy. Nuclei (DAPI) are shown in grey, actin (phalloidin) in red, microvilli (anti-villin) in yellow, tight junctions (anti-occludin) in dark blue, Colonocytes (anti-KRT20) in cyan, Goblet cells (anti-muc2) in green, and enteroendocrine cells (anti-ChrA) in magenta. Main panels, XY projection; right panels, YZ projection; bottom panels, XZ projection. F) Co-culture experimental design with *P. micra* ATCC 33270 (*PmA)*, HHM BlNa17 (*PmB)*, and *F. magna*. G) Cell viability after 48 hours of co-culture under hypoxic conditions at 37°C in 5% CO_2_. Living cells were quantified by flow cytometry and presented as the ratio of living cells in stimulated versus non-stimulated samples (NS). H) Quantification of NF-κB positive nuclei upon immunofluorescent labeling with an anti-p65 antibody and nuclear staining. Results are expressed as ratios of p65-positive nuclei over the total number of nuclei and normalized by the NS condition I) Global DNA methylation dosage by quantification of 5-methyl-cytosine (5m-C) residues compared to total cytosines, after 48 hours of co-culture of bacteria with human colonics monolayers. Results are expressed as the percentages of 5-mC normalized to the non-stimulated samples (NS). In G, H, and I, results are from cells derived from two donors with at least 3 independent experiments per donor and the data are represented as mean +/- SEM. Mann-Whitney test ***p<0.001, **p<0.01, *p<0.05. See also Figure S2. J-K) Gene number and distribution of hypo- and hyper-methylated genes between *P. micra* ATCC 33270 (*PmA*), *P. micra* HHM BlNA17 (*PmB*) phylotypes, or *F. magna* ATCC 29328 and the NS condition. Differentially methylated genes were obtained by calculating a combined p-value based on differentially methylated CpG sites located at specific genomic regions (TSS, in gene promoters; or non-TSS, not at promoter sites) and regions within CpG islands or not. L) Venn diagram of differentially methylated genes when only promoters (TSS) and CpG islands were considered. M, N, O) Volcano plots of differentially methylated genes (hypo- and hyper- methylated) in the comparison of *PmA, PmB* or *F. magna* bacterium- *versus* non-stimulated condition, respectively. CpG sites only in promoters (TSS) and in CpG islands were considered. Genes belonged to oncogenes, tumor suppressor genes (TSGs) or epithelial-mesenchymal transition (EMT) databases, are highlighted by their names. See also Table S3.

*P. micra* has been described as an anaerobic microorganism^28^ and little is known about its oxygen tolerance. To determine compatible oxygenation conditions with eukaryotic cell growth requirements, *P. micra* was cultivated at various oxygen concentrations (0, 2, or 21% O_2_) and viability was assessed by Colony Forming Units (CFU) counting at different time points. *P. micra’s* viability dropped by 70% in less than ten minutes in aerobic conditions (21% O_2_), whereas in hypoxic conditions (2% O_2_) it was able to grow albeit at half the rate compared to full anaerobic conditions (Figure S2A). Thus, primary cells were grown for four days in aerobic conditions to allow monolayer formation, followed by three days in hypoxia (2% O_2_). The primary cell monolayer’s integrity was maintained in these conditions (Figures 3C and D) and showed all characteristics of a differentiated epithelium as assessed by the presence of tight junctions, differentiated epithelial cells, colonocytes, goblet cells, and enteroendocrine cells (Figure 3E). To investigate the possible involvement of *P. micra* in early oncogenic processes, we co-cultivated primary cell monolayers with the two reference isolates, *PmA* and *PmB,* as well as *F. magna* as a control, which is the closest phylogenetic taxon to *P. micra*. One day prior to co-culture, cells were placed in hypoxic conditions (2% O_2_) to allow pre-adaptation. On the fourth day of the culture, cells were co-cultured with bacteria using a multiplicity of infection (M.O.I.) of 250 and incubated at 2% hypoxic conditions for 48 hours prior to analysis (Figure 3F). Bacterial growth was monitored over time by CFU counting of co-culture supernatants. After 48 hours of co-culture, *PmA* showed a 5-fold increase but did not impact cell viability, while *PmB* and *F. magna* showed no significant growth but had a more toxic effect, inducing a 30% decrease in cell viability (Figure 3G). Cell proliferation was measured by Ki-67 quantification upon immunofluorescent staining and no significant differences were observed between the various groups (Figure S2C, F). Because goblet cells are known to be impacted by intestinal microbes^29^, we quantified their proportion and no significant differences were observed (Figure S2D, G). Moreover, because virulent bacteria can stimulate NF-κB and induce DNA breaks ^30^ the proportion of cells with DNA double-strand breaks was measured and was found unchanged (Figure S2E, H). Yet, as compared to the NS condition, a significant increase in nuclear NF-κB was observed with both *PmA* (p<0.01) and *PmB* (p<0.05), but not *F. magna*, (Figure 3H). Thus, in the time frame of the experiments, *P. micra* did not induce DNA breaks while triggering activation of the NF-κB master regulator of inflammatory and anti-apoptotic responses. These results suggested that *P. micra* might contribute to host epithelial cell transformation through a NF-κB pathway-mediated effect.

As we previously showed a correlation between *Parvimonas* carriage in feces and a positive CMI in patients^13^, global DNA methylation was measured using a 5-methyl-cytosine dosage assay *in vitro*. Co-cultures of both *P. micra* phylotypes showed a significant increase in global DNA methylation of primary cells as compared to NS condition (p<0.05) or to *F. magna* (Figure 3I). To identify affected genes, a genome-wide DNA methylation analysis was performed on the human colonic primary cells exposed or not to *P. micra* phylotypes or *F. magna*, using the Infinium MethylationEPIC BeadChip from Illumina. The methylation status of >850,000 CpG sites was assessed (Data set ready on Mendeley Data with the following DOI: 10.17632/nwj3bgbg5m.1). Distribution of CpG sites were then classified by context (in or outside of CpG islands) and regions: body genes (3’ or 5’ UTR, first exon, exon bond, and body), promoters (200 or 1500 bases upstream of the transcriptional start site [TSS200 and TSS1500]) or intergenic. Using the LIMMA paired test we identified hyper- and hypo-methylated CpG sites from colonic primary cells challenged with bacteria compared to non-stimulated cells. CpG sites were then sorted into four genomic categories: CpGs in promoters and within CpG islands; CpGs in promoters and outside CpG islands; CpGs in body genes (promoters region excluded) and within CpG islands; as well as CpGs in body genes outside of CpG islands. Because aberrant DNA methylation in promoters of tumor suppressor genes (TSGs) is a hallmark of CRC, combined p-values of differentially methylated CpG sites within the same gene were determined based on these four categories. We found that DNA methylation modifications were induced by all three bacteria while *PmB* promoted the highest number of differentially methylated genes (DMGs) (Figure 3J). Focusing on CpGs located in promoters and within CpG islands, we identified 15, 90, and 39 DMGs between non-stimulated cells and cells co-cultivated with *PmA*, *PmB*, and *F. magna,* respectively. Venn diagram revealed six genes in common between *PmB* and *F. magna*, only one in-between the two phylotypes of *P. micra,* and two genes for *PmA* and *F. magna* (Figure 3L). Strikingly, most of the differentially-methylated promotors in *PmA* co-culture cells were either found in oncogenes, TSGs, or genes involved in epithelial-mesenchymal transition (EMT) when compared to *PmB* or *F. magna* co-cultures (Figure 3M, 3N, 3O). Cells co-cultivated with *PmA* presented hyper-methylation of several TSGs promoters such as SCIN, HACE1, TSPAN13, FBXO32, IGFBP7, SIX1 or CXXC5. Except for the KIAA0494 gene that codes for an uncharacterized protein, all DMGs induced by *PmA* were involved in carcinogenesis, particularly in EMT processes or cytoskeleton remodeling (Table S3). These results suggest that *P. micra* phylotype A might altered expression of a set of genes, including well-characterized carcinogenesis regulators, through epigenetic promoter gene methylation.

### *P. micra* phylotype A carriage in CRC patients is associated with DNA methylation modifications and gene expression variations

To confirm that the differences in phylotypes might influence colon cancer progression, we put our efforts in isolating *Parvimonas* from colonic tumoral biopsies. We eventually successfully isolated one clone that we could cultivate *in vitro*. The sequencing of the full-length 16S rDNA (Figure 4A) as well the phenotypic properties (eg. haemolysis, density of colonies and adhesion), indicated that this new colonic tumoral isolate, named *PmG5*, belongs to phylotype A. Indeed, *PmG5* was hemolytic, “non-compact” on blood agar plates, and able to adhere to Matrigel® (Figure 4B) and to the colonic cell line TC7 *in vitro* (Figure 4C). No spiky structures were observed on *PmG5* surface by TEM (Figure 4D). Thus, *PmG5* showed all the features of the *PmA* isolate we used for the *in vitro* assay.

**Figure 4.**
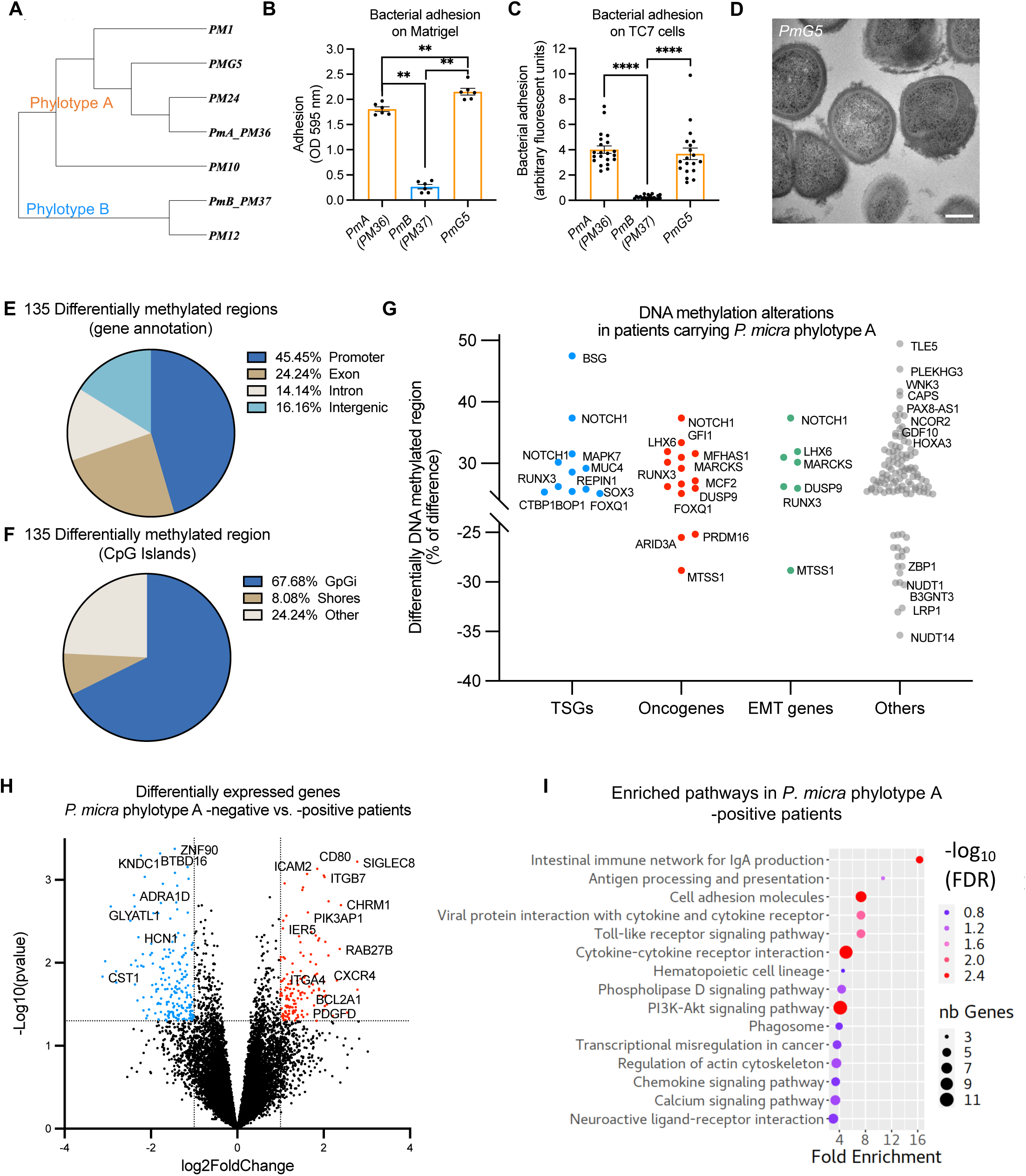
*P. micra phylotype A* carriage in CRC patients is associated with DNA methylation modifications and gene expression variations. A) Phylogenetic tree based on full-length 16S rDNA sequences of *P. micra PmA, PmB*, other oral clinical isolates and of the new isolate from tumoral colonic tissue *PmG5*, showing that it belongs to phylotype A. B) Adhesion capacity of *PmA, PmB and PmG5* to Matrigel®. Optical density at 595 nm represents measurement of bacterial adhesion. C) Adhesion capacity of *PmA*, *PmB* and *PmG5* to human colonic cell lines TC7 after 1 hour of co-culture. Bacterial adhesion was quantified by the analysis of fluorescent images of cells co-cultured with the two phylotypes and is reported as arbitrary fluorescent units. Mann- Whitney test ****p<0.0001; **p<0.01. D) Transmission electron microscopy on ultrafine sections of *PmG5* (Scale: 200 nm. E) Genomic localization and F) CpG context of the 135 differentially methylated regions between three *Parvimonas*-negative and three *Parvimonas*-positive CRC patients, by using reduced-representation bisulfite sequencing (RRBS). G) Identity of genes with differentially methylated regions between *P. micra* phylotype A -negative versus – positive CRC patients. Genes belonged to oncogenes, tumor suppressor genes (TSGs) or epithelial-mesenchymal transition (EMT) databases, are highlighted in red, blue and green respectively. H) Volcano plot of differentially expressed genes (DEGs) between *P. micra* phylotype A -negative versus – positive CRC patients. Blue and red dots represent significantly down and up-expressed genes, respectively (p<0.05; êlog2FoldChangeê>1). H) The Kyoto Encyclopedia of Genes and Genomes (KEGG) pathway enrichment analysis for up-regulated genes in *P. micra* phylotype A positive CRC patients. The x-axis indicates the degree of KEGG pathway enrichment. The y-axis indicates the name of the KEGG pathway. The dot size means the gene number. The dot color indicates the value of the False Discovery Rate (FDR). See also Tables S4.

Having shown that *P. micra* phylotype A is more prevalent in CRC and associated to cancer tissue, we wondered if its presence was also associated to DNA methylation changes in host colonic cells, as observed *in vitro*. We thus performed reduced-representation bisulfite sequencing (RRBS) analysis on colonic tumoral tissues either negative or highly colonized by *P. micra* from 3 patients in each group. We observed 135 differentially methylated genomic regions (DMRs), mostly located in promotor regions (45%) and in CpG island context (68%) (Figures 4E, 4F). Albeit we did not identify the same genes as in our *in vitro* study, differentially methylated regions were found in numerous TSGs, oncogenes, and genes involved in EMT (Figure 4G, Tables S4).

To identify altered gene expression patterns linked with *P. micra* colonization of the mucosa, RNA-seq was performed on colonic tumoral tissues, and transcriptome comparisons of either *Parvimonas*-positives (n=30) or *P. micra* phylotype A-positive patients (n=20) versus negative patients (n=12) were analyzed. We found 483 and 336 differentially expressed genes (DEGs) for *Parvimonas* and *P. micra* phylotype A, respectively (Log2FC>1, p<0.05) (Figure 4H, Tables S4). To uncover potential signaling pathways that are impacted by *P. micra* colonization, we performed Kyoto Encyclopedia of Genes and Genomes (KEGG) pathway analysis (Figure 4I) on over-expressed genes in *P. micra* positive patients. Over-expressed genes in *P. micra* phylotype A -positive patients revealed significant enrichment in several pathways related to inflammation, such as phagosome, Toll-like receptor signaling pathway, cytokine-cytokine receptor interaction, chemokine signaling pathway or intestinal immune network for IgA production, the latter demonstrating a localized inflammatory response to mucosal colonization (Figure 4I). Affected pathways also included transcriptional misregulation in cancer, hematopoietic cell lineage, and PI3K-Akt signaling pathway, a pathway well-known to be involved in cell growth, survival, cell-cycle progression, and differentiation. In addition, DEGs enriched in *P. micra* phylotype A -positive patients were involved in cell adhesion and regulation of actin cytoskeleton (Figure 4I, Tables S4).

These findings revealed that tissues colonized with *P. micra* phylotype A display altered DNA methylation patterns and expression in keys genes involved in carcinogenic events.

## Discussion

Recently, we and others observed an epidemiological association between CRC and several oral bacteria detected in feces using 16S rDNA sequencing^24, 31^ or metagenome analysis^12, 13, 32^. They mainly belong to species such as *Fusobacterium nucleatum*, *Porphyromonas gingivalis*, *Solobacterium moorei*, *Peptostreptococcus stomatis*, *Gemella morbillorum,* and *Parvimonas micra*. Some of these oral bacteria were also associated with colonic tissues of CRC patients^24, 33, 34^, being more abundant in tumors than in normal adjacent tissues^25, 34^. Interestingly, we found that these bacteria were often found in co-occurrence in the mucosa of patients (data not shown), suggesting that these oral microorganisms could live in close community in association with the colonic mucosa, partially reproducing an ecosystem similar to the oral cavity. Indeed, *P. micra, F. nucleatum, P. stomatis,* and *G. morbilorum* have been observed in biofilm-like structures at the surface of the colonic mucosa of CRC patients and healthy subjects^11, 35–39^. In the present study, we particularly focused on *P. micra,* at the subspecies level, and quantitatively measured its carriage in both feces and colonic tissues in a large cohort from Henri Mondor Hospital – Assistance Publique des Hôpitaux de Paris (Directed by Prof. I. Sobhani) that includes normal colonoscopy, adenomatous and CRC patients. We observed, as others^40^, that *P. micra* is present but not significantly enriched in patients with adenomas, these benign tumors being considered as precursor of CRC^41^, indicating that *P. micra* is more likely to be involved in accelerating and/or exacerbating the carcinogenic processes rather than being a primary driver bacterium, according to the “driver-passenger” model^42^. And indeed, genetically predisposed murine model of intestinal adenoma formation (e.g., Apc^Min/+^ mice) orally gavaged with *P. micra* exhibited a significantly higher tumor burden and were associated with altered immune responses and enhanced inflammation^43, 44^. In human, *P. micra*, like other oral pathobionts, have been shown to be associated with the CMS1 subtype of CRC, which makes up 14% of all CRC cases^45^, and with an over-activation of genes involved in immune responses. The CMS1 tumors, also called “immune subtype”, are characterized by a strong immune cell infiltration by CD8+ cytotoxic T cells, CD4+ T helper cells, and natural killer cells^46, 47^. Therefore, a key aspect of *P. micra* involvement in colon carcinogenesis is inflammation.

Next steps require moving from association to causality to identify the mechanisms and define more aggressive phylotypes driving carcinogenesis. In the present study, we have identified different phylotypes of *P. micra*, observed a strong association of the adherent and hemolytic phylotype A with CRC, and developed a physiologically relevant low-oxygen *in vitro* co-culture model, to assess the interaction of this particular pathobiont with primary human colonic intestinal cells. After 48h of co-culture, no impact of *P. micra* was observed on cell proliferation, differentiation, or DNA damage. In contrast, *P. micra* induced the activation of the central transcription factor NF-κB, a master regulator of inflammation and anti-apoptotic responses, involved in gene-expression reprogramming in cancers. In a recent *in vitro* study, tumoral colonic cell lines were infected with *P. micra* and presented a higher proliferation rate upon infection due to the activation of the Ras signaling pathway^44^. Consistent with this observation, we showed that patients carrying *P. micra*, have altered PI3K-Akt signaling, a downstream effector of the Ras pathway, demonstrating that *P. micra* phylotype A might induce aberrant cellular proliferation that may contribute to initiation and progression of cancer cells.

Apart from specific somatic mutations that characterize cancerous cells, epigenetic DNA modifications, particularly methylation, of TSGs promotors are main contributors in colonic carcinogenesis. We recently showed that CRC-associated dysbiotic feces transplanted to mice caused epigenetic changes similar to those observed in human tumors and the occurrence of murine colonic crypt aberrations^13^. Furthermore, the CMS1 CRC subtype, in which *P. micra* and other oral bacteria enrichment has been observed^45^, is associated with the phenotype of CpG islands hypermethylation in TSG promoters (high CIMP)^46^. Consistent with this hypothesis, Xia *et al.* observed an association between enrichments of *F. nucleatum*, *Parvimonas spp.* or other bacteria, and hypermethylation of promoter in several TSGs in CRC tumoral tissue. Moreover, they observed that *F. nucleatum* was able to up-regulate DNA methyltransferase activity *in vitro*^48^, although they did not focus on *Parvimonas* subspecies. Further, we have shown in a large CRC patients’ cohort that *P. micra* was significantly associated with methylation of the *WIF1* promoter, a very well-known TSG^13^. In the present study, we report for the first time that *P. micra* increases global DNA methylation of target host cells using an *in vitro* co-culture model of human primary colonic cells. Notably, by comparing different *P. micra* phylotypes, we established a *P. micra* phylotype A-dependent signature of methylated promoters whose gene functions converge towards the regulation of the cytoskeleton (Table S3) and include TSGs or genes involved in EMT (Table S3). For example, Scinderin gene (*SCIN*) coding for a Ca^2+^dependent actin-severing and capping protein, is involved in the regulation of actin cytoskeleton and known to be overexpressed in CRC^49^; Tetraspanin 13 (*TSPAN13*) is a TSG coding for a transmembrane signal transduction protein that regulates cell development, motility, and invasion^50^; *DIAPH3* gene, is a major regulator of actin cytoskeleton involved in cell motility and adhesion^51^; Semaphorin 3F (*SEMA3F*), is a TSG coding a secreted protein involved in cytoskeletal collapse and loss of migration^52^; and *SASH1* is a TSG coding for a scaffold protein involved in the TLR4 signaling, and known to interact with the actin cytoskeleton to maintain stable cell-cell adhesion^53^. Notably, the hypermethylation of the TSG *HACE1* gene promoter was also observed upon co-culture with *P. micra* phylotype A. HACE1 is an E3 ubiquitin ligase controlling RAC1 stability, a small GTPase of the Ras superfamily involved in cell motility^54^ and the TSG *HACE1*’s promoter was previously shown to be hypermethylated in CRC^55^. Hence, phylotype A of *P. micra* could contribute at several stages of CRC development by causing DNA methylation modifications enhancing cell transformation through cytoskeleton rearrangement. By comparing the methylome and transcriptome of *P. micra*-negative versus-positive patients, we observed differentially methylated regions in several other TSGs, oncogenes or genes involved in EMT, together with significant enrichment of DEGs involved in cell adhesion, regulation of cytoskeleton, and PI3K-Akt signaling that promotes cell-cycle progression, cell survival, migration, and tumor progression.

In conclusion, here we described the *P. micra* phylotype A as the most prevalent CRC-associated *Parvimonas*, characterized by its hemolytic capacity and adherent properties. This pathobiont colonizes the colonic mucosa, induces inflammation, and causes DNA methylation changes, creating a proper ground for carcinogenesis by potentially allowing cells to accumulate supplementary genomic alterations. *P. micra*, together with some other similar pathobionts of the oral cavity, by modifying epigenetic marks on the genome, would promote further transformations, comprising enhancement of cell migration, invasion, or metastatic events, that could impact tumor aggressiveness. Clearly, further investigation is warranted on the cross-talks among and between pathobionts, including *P. micra*, and the human colonic epithelium. As an example, bacterial hemolysins have been shown to cause epigenetic marks in histones^56^ but studies on DNA methylation changes are lacking.

## Acknowledgments

The authors thank patients for having contributed to the present research program. We thank the bacteriology Departments of Cochin, Pitié Salpétrière, and Mondor Hospitals from APHP for providing *P. micra* clinical isolates (particularly Drs Asmaa Tazi, Alexandra Aubry and Biba Nebbad) and the Collection of Institut Pasteur for providing reference strains. We acknowledge the Service de Gastroenterology (Dr M. Uzzan) and URC (F. Canoui Poitrine) of Henri Mondor, and the URC of St Antoine hospitals at APHP (Prof. T. Simon), as well as Center for Translational Science (CRT)-Cytometry and Biomarkers Unit of Technology and Service (CB UTechS) at Institut Pasteur for their supports. We are grateful for support for UBI equipment from the French Government Programme Investissements d’Avenir France BioImaging (FBI, N° ANR-10-INSB-04-01) and the French government (Agence Nationale de la Recherche) Investissement d’Avenir programme, Laboratoire d’Excellence “Integrative Biology of Emerging Infectious Diseases” (ANR-10-LABX-62-IBEID). We thank Claude Parsot and Shaynoor Dramsi for their dedication and inspiration in the development of this work, and Pamela Schnupf for careful reading of the manuscript. This study was performed with the financial support from the Université Paris Est Creteil (UPEC), PHRC (Vatnimad) of the French government, SNFGE (French society of Gastroenterology Commad Support 2019), and from ITMO Cancer AVIESAN (Alliance Nationale pour les Sciences de la Vie et de la Santé, National Alliance for Life Sciences & Health) within the framework of the Cancer Plan (HTE201601) supported by INCA, and executed within the frame of the Oncomix research program between Institut Pasteur and AP-HP.

## Authors contributions

Clinical and translational concept IS; basic conceptualization of *in vitro* study PS, GN, and EB; methodology GN, EB, AM, EL; investigation CB, FD, TP, GN and EB; formal analysis AD, DM, SV, AV, GN and EB; writing – original draft GN and EB; writing – review & editing GN, PS and IS; funding acquisition and resources PS, MC, IS AM, and EL and supervision PS, IS and GN.

## Materials and Methods

### Patient recruitment at the Créteil Henri Mondor Hospital

Participants were selected from different cohorts recruited with informed consent between 2004 and 2018 by the endoscopy department at Henri Mondor hospital (Créteil) where patients had been referred for colonoscopy; detailed description in ^57^. Participants with previous colon or rectal surgery, colorectal cancer, inflammatory bowel diseases, or with a genetic form of CRC were excluded. Individuals exposed to antibiotics or probiotics within four weeks before collection or suffering from acute diarrhea were also excluded from the study. Tumor Node Metastasis (TNM) stages of colonic neoplasia were determined by radiological examinations and analyses of the surgical specimen by the anatomopathological department of Henri Mondor Hospital. Clinical parameters of the patients, such as body mass index (BMI), age, sex, and disease history were referenced. The cumulative methylation index (CMI) score was previously determined ^57^. This score was calculated from the methylation status of three genes (*wif1*, *penk* and *<u>npy</u>*) involved in colorectal carcinogenesis and was considered negative for CMI<2 or positive for CMI≥2. Clinical data are summarized in Table S2.

### Fecal and tissue samples

Fresh feces were collected between 2 weeks and 3 days prior to colonoscopy and ten grams were frozen at −20°C for 4 hours and then stored at −80°C until use. Paired samples of colorectal tumor tissue and homologous normal mucosa (more than 15 cm from the margin of tumor resection) were collected within 30 minutes after surgical resection and immediately frozen in liquid nitrogen and stored at −80°C until further use.

### DNA extraction and quantification

Fecal DNA extraction was performed using the G’NOME DNA isolation kit® (MP Biomedicals) according to the supplier’s instructions, with modifications as described in ^58^. Tissue DNA extraction was performed from eight 50 μm cryosections of nitrogen frozen tissue using the QIAamp PowerFecal DNA Kit® (Qiagen) following the manufacturer’s instructions with the following modification: 0.1 mm diameter silica beads were added to the lysis solution provided and shaking was performed at maximum speed for 10 minutes in a vibratory shaker. Fecal and tissue DNA concentrations were determined by Qubit® fluorometer (ThermoFisher) and stored at −20°C until use.

### Whole genome sequencing (WGS) on fecal samples

Metagenome sequencing was performed by the high-throughput platforms BIOMICS (Institut Pasteur, Paris, France) and EMBL (Heidelberg, Germany). Sequencing was performed in pairs, using the HiSeq 2000/platform 2500 equipment, over a length of 100 bp of DNA and at a sequencing depth of 5 Gbp ^26^. The raw data have already been reported in several papers ^12, 26, 57^. For this study, one hundred and sixty-six fecal samples from the CCR1 and DETECT cohorts were considered. The Diamond/MEGAN6 bioinformatics pipeline ^59^ was used for metagenomic assignment. Sequences were filtered for an average quality (Phred score) greater than 20 over a window of two consecutive bases and a length greater than 100 bp using Trimmomatic software (version 0.35). Good quality sequences were translated in the six possible reading frames and aligned to the NR reference library (RefSEQ non-redundant protein database) in which proteins/peptides that are more than 99% similar are combined into a single organism-associated group with a specific identifier ^60^. A sequence can be assigned to several taxa. The MEGAN6 LCA (Lowest Common Ancestor) algorithm ^61^ was used to resolve multi-mapping reads: when a sequence was assigned to two (or more) taxa of different phylogeny, it was assigned to the top taxonomic level in common.

### 16S rDNA sequencing of tissue samples

For this study, 71 pairs of homologous tissue samples (matched tumor and healthy tissue) were analyzed. After amplification of the 16S rRNA gene V3-V4 region, pairwise sequencing was performed to a length of 250 bp on the Illumina MiSeq platform. The resulting raw database was cleaned of sequences corresponding to human or phage sequences. Adapters and primers, as well as 5’ and 3’ ends with a Phred quality score over a 2 bp sliding window of less than 30 were removed using Trimmomatic (version 0.35) software. Paired sequences were merged using the FLASH2 tool (version 2.2.00). Taxonomic assignments were performed using MALT/MEGAN6 bioinformatic pipeline ^61^ with default parameters and the SILVA rRNA database version 123 ^62^.

### RNA sequencing and pathway enrichment analysis

Total RNA was extracted from tumoral colonic mucosa samples using TRIzol®-chloroform extraction method. RNA was sent to the NOVOgene company (China) that carried out the quality control, library preparation, and sequencing. The reads quality was assessed using FastQC (v0.11.4 https://www.bioinformatics.babraham.ac.uk/projects/fastqc/) combined with MultiQC (v1.6 http://multiqc.info/) ^63^. We performed splice-aware alignment on RNAseq using the STAR transcriptome aligner (v2.5.0) ^64^ with human genome version GRCh38 from Ensembl release 99 (Homo_sapiens.GRCh38.dna.primary_assembly.fa). After alignment, featureCounts (subread v 1.6.1) ^65^ was used to obtain the matrix of counts of fragment by genes from sorted BAM files. We removed samples for which less than 50% of the fragments mapped to genes and used blast+ (2.9.0) to align the reads on the nt collection and confirm that the lack of alignment to human genes was due to an excess of reads mapping to non-human organism, suggesting a contamination. Overall, 88 samples were retained after quality control. We filtered gene level raw counts by removing all genes with no fragment mapped in more than half of the samples. Normalized counts for further use were generated by applying a quantile normalization using the voom procedure ^66^ implemented in the limma package (v3.46.0) ^67^. The 55 tumor tissue samples from which we had information both for *Parvimonas* carriage, assessed by 16S rDNA sequencing, and host gene expression, were kept for differential expression analysis. Patients with no matching reads (dp=0) in neither *Parvimonas genus*, *P. micra* phylotype A or *P. micra* B sequences, were considered as negative (n=12). Patients with at least 10 matching reads in *Parvimonas* sequence were considered as *Parvimonas*-positive (n=30). Patients with at least 10 matching reads in *P. micra* phylotype A sequence and representing at least 70% of all *Parvimonas* reads were considered as *P. micra* phylotype A-positive patients (n=20). Differential analysis was not performed for the phylotype B as only 3 patients were *P. micra* phylotype B-positive. The linear modeling followed by empirical Bayes statistics approach implemented in limma ^68^ was used to assess differential expression. Differentially expressed genes (DEGs) were identified as p < 0.05 and log2 fold change (Log2FC) > 1, and subjected to functional classification analysis, using ShinyGO (version 0.77) ^69, 70^ and Kyoto Encyclopedia of Genes and Genomes (KEGG) database ^71^. Pathway enrichment results were double check using DAVID (Database for Annotation, Visualization, and Integrated Discovery) version 2021 ^72, 73^.

### Colonic cell cultures

Cancer cell lines: human colon cell lines HT-29, HCT116 and TC7 (subclone of Caco-2), were grown in DMEM (“Dulbecco Modified Eagle Medium”) 1 g/L glucose (Gibco), supplemented with 20% (vol/vol) decomplemented fetal bovine serum (FBS) (Gibco), 1% non-essential amino acids (Gibco), 1% GlutaMAX (Gibco), at 37°C in 10% CO_2_.

### Human colonic organoids

Human colonic surgical resection specimens were obtained from Henri Mondor Hospital from two patients who had undergone colectomy surgery from rectal adenocarcinoma and had given their informed consent (agreement N°2012-37). Tissues samples from the normal site (far from tumor tissue) were sterilely washed using PBS supplemented with gentamicin (50 µg/mL), Normoxin (1 mg/mL) and amphotericin B (2 µg/mL), to obtain bacteria-free normal colonic mucosa and kept in 0.1% PBS-BSA at 4°C for the duration of transport (approximately 4 hours). Crypts were isolated according to the protocol of Sato et al, (Sato et al., 2011), with some modifications. The epithelium was stripped of underlying muscularis and serosa, washed several times in cold PBS until the supernatant was clear, and cut into 5 mm^2^ fragments. The fragments were incubated for 20 minutes on ice in cold chelation buffer consisting of 5.6 mmol/L Na_2_HPO_4_, 8.0 mmol/L KH_2_PO_4_, 96 mmol/L NaCl, 1.6 mmol/L KCl, 44 mmol/L sucrose, 54.8 mmol/L D-sorbitol, and 0.5 mmol/L DL-dithiothreitol in distilled water plus 2 mmol/L EDTA. After removal of the supernatant and addition of cold chelation buffer without EDTA, the tissue fragments were vigorously resuspended by several passes through a 10 mL serological pipette. After sedimentation of the tissue fragments, the supernatant, containing the crypts, was recovered. The resuspension/sedimentation procedure was repeated twice. The tissue fragments were again incubated in chelation buffer with EDTA for 15 minutes and the resuspension/sedimentation procedure was repeated three times. Fractions containing crypts were pooled and centrifuged at 300g for 5 minutes. The crypt pellet was resuspended in Matrigel® growth factor-reduced medium (Corning) diluted to 75% in culture medium (see composition below). Four 25 µl drops of the Matrigel-crypt mixture were placed per well in 12-well plates with approximately 100 crypts per drop. The plates were incubated for 15 minutes at 37°C to allow polymerization of the Matrigel, then 800 μL of culture medium was added and the plates were incubated at 37°C and 5% CO_2_. The culture medium consisted of AdvancedDMEM/F12 (Gibco), 10 mM HEPES (Gibco), 1X GlutaMAX (Gibco), 100 υ/mL penicillin, 100 µg/mL streptomycin (Gibco), 1X N2 (ThermoFisher), 1X B27 (ThermoFisher), 1 mM N-acetyl-L-cysteine (Sigma), 100 ng/mL Noggin (R&D systems), 50 ng/mL recombinant human EGF (R&D systems), 150 ng/mL Wnt-3A (R&D systems), 1 µg/mL recombinant human R-spondin-1 (R&D systems), 500 nM A83-01 (R&D systems), 10 mM nicotinamide (Sigma), 10 µM SB202190 (Sigma), 10 nM Gastrin I (Sigma), 3 µM CHIR99021 (Biogems), 10 µM Y27632 (Sigma) and 10% decomplemented fetal bovine serum (vol/vol) (Gibco). The culture medium was changed every two days, without Y27632. Each week, organoids were split with a 1:3 ratio. The medium was replaced by cold fresh medium and the Matrigel drops with the organoids were dissociated by pipetting several times through a 200 µl tip and collect in a tube. After a spin for 5 minutes at 300g at 4°C, the pellet was washed with fresh cold medium, resuspended with a p200 pipette several times, and spun again. The pellet of organoid fragments was resuspended and diluted in Matrigel® as described above. After expansion, organoids were frozen and stored at −80°C in freezing medium composed of Advanced DMEM/F12, 10% fetal bovine serum, 10% DMSO (PAN Biotech), 10 µM Y-27632 and 3 µM CHIR99021.

### Generation of human colonic epithelial monolayer

Monolayers of organoid-derived primary cells were generated as previously described ^27^. Briefly, the twenty-four well plates with 0.4 µm pores (Corning) Transwell® insert system were used as culture support for primary cells in monolayers. Inserts were previously coated with 50 µl of human collagen IV at 30 µg/ml (Millipore) overnight at 4°C, then washed with DMEM. Approximately three hundred organoids at five days of growth were dissociated into fragments (using Cell Recovery (Corning)), resuspended in 200 µl of culture medium and put on each insert. The culture medium consisted of DMEM/F12 (Gibco), 10 mM HEPES (Gibco), 1X GlutaMAX (Gibco), 100 u/mL penicillin, 100 µg/mL streptomycin (Gibco), 1X N2 (ThermoFisher) and B27 (ThermoFisher) supplements, 1 mM N-acetyl-L-cysteine (Sigma), 100 ng/mL noggin (R&D systems), 50 ng/mL recombinant human EGF (R&D systems), 150 ng/mL Wnt-3A (R&D systems), 1 µg/mL recombinant human R-spondin-1 (R&D systems), 500 nM A83-01 (R&D systems), 10 mM nicotinamide (Sigma), 10 µM SB202190 (Sigma), 10 nM Gastrin I (Sigma), 3 µM CHIR99021 (Biogems), 10 µM Y27632 (Sigma) and 10% decomplemented fetal bovine serum (vol/vol) (Gibco). Monolayers were incubated at 37°C in 5% CO_2_. After three days of culture, the medium was changed without addition of Y-27632. Cell growth and monolayer closure were monitored daily by observation with an inverted brightfield microscope (IX81, Olympus) and by recording the transepithelial electrical resistance (Millipore).

### Bacterial strains, culture and characterization

*Escherichia coli* pks+ strain IHE3034 was used as a positive control for double-strand DNA breaks. *Shigella flexneri* 5a BS176, lacking the virulence plasmid, was used as a negative control for bacterial adhesion and autoaggregation negative control. The reference strains ATCC 33270 of *P. micra (PmA)* and ATCC 29328 of *Finegoldia magna* were obtained from the Pasteur Institute Collection (CIP). In this study, twenty-seven clinical isolates of *P. micra*, including HHM BlNa17 (*PmB*), were collected from the bacteriology departments of three Parisian hospitals: Henri Mondor, Cochin, and Pitié Salpetrière (Table S1). The identification of the different isolates was confirmed by mass spectrometry and 16S rDNA sequencing analysis (Eurofins Genomics) and sequences were deposited on GenBank (Table S1). 16S rDNA whole sequence from positions 112 to 1302 were aligned with the ClustalW program (EMBL) and the phylogenetic tree was calculated using the PhyML program. Pan-genomic comparison: available complete *P. micra* genomes were downloaded from the NCBI database. Analysis of pan-genomes and comparison of common or strain-specific gene carriage was performed using the Roary program^74^ with a 95% identity threshold and the use of default parameters. Roary outputs a multi-FASTA alignment of all of the core genes and a matrix of presence or absence of each gene in each genome. The maximum Likelihood phylogenetic tree from the alignment of all of the core genes was inferred using Fastree with the GTR+CAT model. An additional script called roary_plots.py was used to visualize the tree against the presence and absence of core and accessory genes.

### Culture

Bacteria were grown under anaerobic conditions in TGY-V enriched broth (Trypticase peptone 30 g/L, D-glucose 10 g/L, Yeast extract 20 g/L, L-cysteine-HCl 0.5 g/L, haemin 5 mg/L, vitamin B12 5 μg/L, menadione sodium bisulfite 500 μg/L, Thiamine 1 mg/L, nicotinic acid 1 mg/L, riboflavin 500 μg/L, p-aminobenzoic acid 100 μg/L, biotin 25 mg/L, calcium pantothenate 1 mg/L, pyridoxamine 2HCl 500 μg/L, folic acid 500 μg/L) or on Columbia agar solid support enriched with 5% horse blood (COH) (Biorad). Anoxic conditions were generated in Gaspack (BD) or anaerobic cabinet (Don Whitley DG250 Workstation). Hemolysis ability and colonies compactions were assessed by observing CFUs on COH plates after 48 hours at 37°C in anaerobic conditions: i) contact hemolytic activity was defined by the presence of an hemolytic zone located underneath the colony and slightly beyond (Figure S1A) and ii) colonies compaction was defined as “non-compact” or “compact”: the “non-compact” colonies spread out on agar upon contact with a loop, whereas the compact colonies remained associated in clumps suggesting a strong interbacterial aggregative property. For the sedimentation assay, bacteria were grown on blood agar plates under anaerobic conditions at 37°C for 48 hours, resuspended in PBS, and incubated at 4°C statically. The optical density (600 nm) in the upper half of the test tubes was measured periodically (Figure S1B).

### *P. micra* isolation from tumoral colonic biopsy

Biopsies of tumoral colonic mucosa from colorectal cancer patients were recovered at Henri Mondor Hospital. After colonoscopy, biopsies were immediately placed in 0.5 mL Shaedler liquid media and maintained at room temperature and in anaerobic condition for transport. The same day, biopsies were placed inside an anaerobic cabinet (0% of O_2_) where all steps of the isolation procedure were achieved. Biopsies were transferred in 0.5 mL of Shaedler media complemented with EDTA-free Protease Inhibitor Cocktail (Sigma) and 0.1% triton (Sigma), containing 0.2-mm-diameter glass beads (EMSCO) and tissue was disrupted upon bead beating agitation carried out at 20/s frequency for 5 min using a Beadbeater (Retsch MM400). Anti-*Parvimonas* serum (on order by Covalab), able to recognize both phylotypes, was add to the mixture at 1:100 and incubated for 30 minutes with agitation. After three PBS washes of tissue fragments, 10^6^ magnetic beads in 0.2 mL PBS (Dynabeads ProteinA, Invitrogen) were added and incubated 15 minutes at room temperature upon agitation. Using a magnet, the beads were washed three times and further incubated for 48 hours at 37°C in TGY-V media. Several dilutions of the growth culture media were plated on blood agar plates and CFUs were identified using PCR (*P. micra*; PF: 5’-GACGGGCGGTGWGTRCA-3’; PR: 5’-AGAGTTTGATCCTGGGTCAG-3’).

### Transmission electron microscopy

Bacteria from a 4-day-old cultures were fixed by adding glutaraldehyde (2.5% final concentration) to the culture for 1 hour at room temperature (RT). The bacteria were washed twice with HEPES 0.1 M, pH 7.5 and postfixed with 1% osmium tetroxide in HEPES 0.1 M pH 7.5 for 1h30 at RT. After three washes in distilled water, the bacteria were dehydrated in a graded ethanol series (25, 50, 75, 95 and 100% ethanol), and gradually infiltrated in Epon resin. Ultrathin sections (60 nm) were obtained on a FC6/UC6 ultramicrotome (Leica). Sections were transferred to 200 Mesh Square Copper grids (CF-200-CuO, Delta Microscopy) formvar and carbon coated, stained with 4% uranyl acetate, and counterstained with Reynold’s lead citrate for 20 minutes. Images were recorded with TECNAI SPIRIT 120 Kv, with a bottom-mounted EAGLE 4KX4K Camera.

### Bacteria adhesion

Adhesion to extracellular matrix proteins: MaxiSorp 96-well flat-bottom plates (ThermoFisher) were coated overnight at 4°C under agitation with 1 μg per well, in triplicate, with the following extracellular matrix proteins: collagen I (Gibco), collagen IV (Millipore), fibronectin (Gibco), fibrinogen (Sigma), laminin (Sigma), and Matrigel® (consisting of a mixture of proteins, mainly laminin, collagen IV, entactin, and heparan sulfate proteoglycan) (Corning). Bacteria were grown on COH plates for 48 hours at 37°C under anaerobic conditions, collected and suspended in PBS at an OD of three units. The bacterial suspension was placed in the wells and the plate was centrifuged at 300g for 5 minutes. After incubation for 1 hour at RT, the wells were washed three times with PBS to remove non-adherent bacteria and then stained with 0.1% crystal violet (Sigma). The plates were incubated for 30 minutes at RT and washed three more times with PBS. The dye was dissolved with 20:80 acetone-ethanol and the optical density at 595 nm was read by a spectrometer (Infinite M200Pro-TECAN).

*P. micra* adhesion to colonic cell lines: human colonic cell lines (TC7, HT-29, and HCT116) were grown on glass coverslips in 24-well plates to confluence. In parallel, *P. micra* was grown on COH plates for 48 hours at 37°C under anaerobic conditions. Bacteria were washed in PBS, re-suspended in DMEM with 1 g/L glucose (Gibco), and put on the cells using a M.O.I. of 1. The plates were centrifuged for 5 minutes at 300g and incubated for 1 hour at 37°C. After incubation, the cells were washed three times in PBS, fixed in 4% paraformaldehyde (PFA) (Electron Microscopy Sciences) for 30 minutes at RT, washed again three times in PBS, and then stored in 0.1% PBS-BSA at 4°C until labeling. Non-specific sites were blocked with 1% PBS-BSA for 1 hour at RT. The cells were then incubated with rabbit anti-*Parvimonas* serum (on order by Covalab) at 1:1000 in 1% PBS-BSA for 30 minutes at room temperature. After washing, the cells were permeabilized with PBS-0.2% triton for 30 minutes. Anti-rabbit secondary antibody coupled to Alexa Fluor 488 (ThermoFisher) at 1:400 and phalloidin coupled to Alexa Fluor 568 (Invitrogen) at 1:200 were used in 1% PBS-BSA for 45 minutes at RT. After washing, nucleic acids were labeled with 1 µg/mL DAPI (ThermoFisher) for 5 minutes. The coverslip with the cells were washed again and mounted on a slide with Prolong Gold (ThermoFisher). Images were taken using a slide scanner with 40X objective (AxioScan, Zeiss) and processed using Fiji ^75^. Bacterial adhesion was quantified as area of the anti-*Parvimonas* fluorescence signal over an entire field normalized by the total number of nuclei and reported as arbitrary fluorescent units x100.

### Co-culture of colonic cells and *P. micra*

Bacteria were grown in TGY-V medium under anaerobic conditions at 37°C for 48 hours, followed by a 1:10 subculture for 24 hours to reach exponential phase. Bacterial density was adjusted to an M.O.I. of 250 in cell culture medium and then put on the cells. Twenty-four hours prior to bacterial exposure, the colonic cells were placed in 2% O_2_ in H35 hypoxic cabinet (Don Whitley) to allow acclimatization. Co-cultures with primary cells were performed for 48 hours at 37°C under hypoxic conditions (2% O_2,_ 5% CO_2_).

### Flow cytometry

Cell monolayers were dissociated with TrypLE™ Express Enzyme (Gibco) at 37°C for ten to twenty minutes. The cells were recovered in DMEM/F12 at RT and then centrifuged for five minutes at 300g. After washing the pellet in PBS, cells were stained with the Live/Dead fixable cell stain kit (ThermoFisher) for twenty minutes on ice, following the supplier’s instructions. Cells were analyzed using a FACS Attune (ThermoFisher) and data were analyzed using FlowJo software (v10.1).

### Ki-67, γH2ax, Muc2, and NF-κB immunofluorescence

Following 48 hours co-culture with *P. micra*, cells were washed with PBS, fixed (4% PFA, 30 minutes), washed again, and stored (0.1% PBS-BSA) at 4°C until labeling. Cells were permeabilized with 0.5% Triton-PBS for 30 minutes at RT, washed in 0.1% PBS-Triton, and blocked with 1% bovine serum albumin (BSA)-0.1% Triton for 30 minutes. The cells were then incubated either i) with a 1:200 dilution of an antibody against the proliferation marker Ki-67 coupled to Alexa Fluor 666 (clone SolA15, eBioscience) and a 1:500 dilution of an antibody against the phosphorylated form of H2A histone family member X (γH2Ax) (clone JBW301, Merck) and 1:200 phalloidin coupled to Alexa Fluor 568; ii) a 1:50 dilution of an antibody against nuclear factor kappa-light-chain-enhancer of activated B cells (NF-κB) p65 (ab7970 Abcam) and 1:200 phalloidin coupled to Alexa Fluor 568; or iii) with a 1:100 dilution of an anti-mucin2 (Muc2) antibody (clone Ccp58, Abcam) and 1:200 phalloidin coupled to Alexa Fluor 568, in 0.1% Triton-BSA overnight at 4°C. After washing in 0.1% PBS-Triton, the cells were incubated with 1:400 secondary antibodies coupled to Alexa Fluor 488 (ThermoFisher) in 0.1% PBS-Triton for 45 minutes at RT. Cells were washed again in 0.1% PBS-Triton and the nucleic acids were labeled with 1 µg/mL DAPI for 5 minutes before cells were mounted on a slide with ProlongGold (ThermoFisher). Fluorescence images were taken using a confocal fluorescence microscope (Zeiss) equipped with an Opterra system (Bruker), and the number of cells was analyzed as the number of positive cells/total cells using Fiji ^75^ or Imaris 9.8 software (URL: http://www.bitplane.com/imaris/imaris).

### DNA methylation studies

Human cells co-cultured with *P. micra* were subjected to DNA extraction using the DNeasy Blood & Tissue Kit® (Qiagen) according to the manufacturer’s instructions and DNA was quantified using a Qubit® fluorometer.

Global DNA methylation level: MethylFlash methylated DNA Quantification Fluorometer Kit (Epigentek) was used to detect the global methylation level in DNA isolated from primary colonic cells co-cultured with *P. micra* ATCC 33270 (*PmA*), HHM BlNa17 (*PmB*) and *F. magna*. Assays were performed in duplicate. As instructed by the manufacturer, 100 ng of the isolated genomic DNA was bound to the assay well. The capture antibody, detection antibody, and enhancer solution were then added consecutively to the wells before the fluoro-developing solution was added and the relative fluorescence units (RFU) were measured (TECAN). The percentage of 5-methylcytosine (5-mC) relative to the total amount of cytosines in the sample was calculated to represent the global methylation dosage and was reported as a ratio to the non-stimulated samples.

DNA methylation profiling: DNA methylation profiling was performed on eight independent co-cultures of *PmA*, *PmB*, and *F. magna* with human primary colonic cells (four experiments per patient tissue). The Infinium MethylationEPIC BeadChip Kit (850K) (Illumina, San Diego, CA, USA) was used for a genome-wide methylation profiling to determine the DNA methylation status of >850,000 CpG sites^76^. A total of 500 ng of genomic DNA from cells in co-cultures were bisulfite-treated using the ZymoResearch EZ DNA Methylation kit (Zymo Research Corp, Irvine, CA, USA). The Infinium HD Methylation Assay (bisulfite modification, amplification, fragmentation, precipitation, hybridization, wash, extension, staining, and imaging) was performed according to the manufacturer’s explicit specifications. The quality was supported by multiple quality control (QC) measures, including tests for proper bisulfite conversion, staining, and specificity of the internal controls, as determined by Illumina GenomeStudio software. Average-beta values (proxy for methylated DNA level between 0, unmethylated, and 1, fully methylated) were normalized to internal controls and corrected by background subtraction. Non-autosomal CpGs (n=14,522) and CpG probes with suboptimal detection (p<0.05 in at 80% of samples) (n= 29,316), as well as single nucleotide polymorphisms (SNPs) associated CpGs (n=59), were removed from our analyses. Beta values were transformed into M-values as described by^77^, and differential probes were assessed using the limma statistical model^78^ package R limma, version 4.0.2. The four conditions (Non-stimulated NS; ATCC; BlNa and FM) and individuals were kept as variables. Pairwise comparisons were obtained from a contrast matrix. No adjustment method was used. For each gene (n=25,560), a t-test on log-fold-change was performed on four categories of probes depending on their CpG context and gene localization: probes in CpG island context and 1/ localized in genes promoters (TSS1500+TSS200) or 2/ in body gene; and probes out of CpG island context 3/ in promoters or 4/ in body gene. Genes with >2probes, p-value <0.05, and with a log-FC >0,1 (absolute value) were considered significant. Using three different databases, the genes were classified as oncogenes, tumor suppressor genes, or involved in Epithelial-Mesenchymal Transition (ONGene^79^, TSGene 2.0^80^ or dbEMT 2.0^81^).

### Reduced Representation Bisulfite Sequencing (RRBS)

Genomic DNA was isolated from Formalin-fixed paraffin-embedded (FFPE) tissue blocks of tumoral colonic tissue, using the Maxwell RSC® DNA FFPE kit (Promega, Madison, WI, USA). DNA concentration was measured with the Qubit® dsDNA HS Assay Kit (Thermo Fisher Scientic, Watham MA), and DNA quality was assessed by gel electrophoresis. Each experimental group (*P. micra* – negative and – positive) contained three different CRC patients. DNA Methylation Profiling (RRBS Service) was performed by Active motif (Carlsbad, CA, USA); DNA digest, adaptor ligation, bisulfite conversion, library amplification, sequencing and data processing. Briefly, Bismark v0.23.0 (https://www.bioinformatics.babraham.ac.uk/projects/bismark/) was used to align reads to a human reference genome (hg38). Methylation analysis was performed with the MethylKit 1.18.0 software^82^. Pairwise comparisons using Fisher’s test were conducted to identify differentially methylated regions (DMRs), consisting of 1000bp regions. SLIM method (sliding window model) was used to correct for multiple comparisons and convert p-values to Adjusted p-value (q-values). Significant DMRs were defined as having a methylation fold change difference greater than 25% and q-value < 0.1. Gene name, genomic region, and location in relation to CpG islands were identified for each DMR.

### Graphical abstract

The graphical abstract was Created with BioRender.com

---

**Figure S1:**
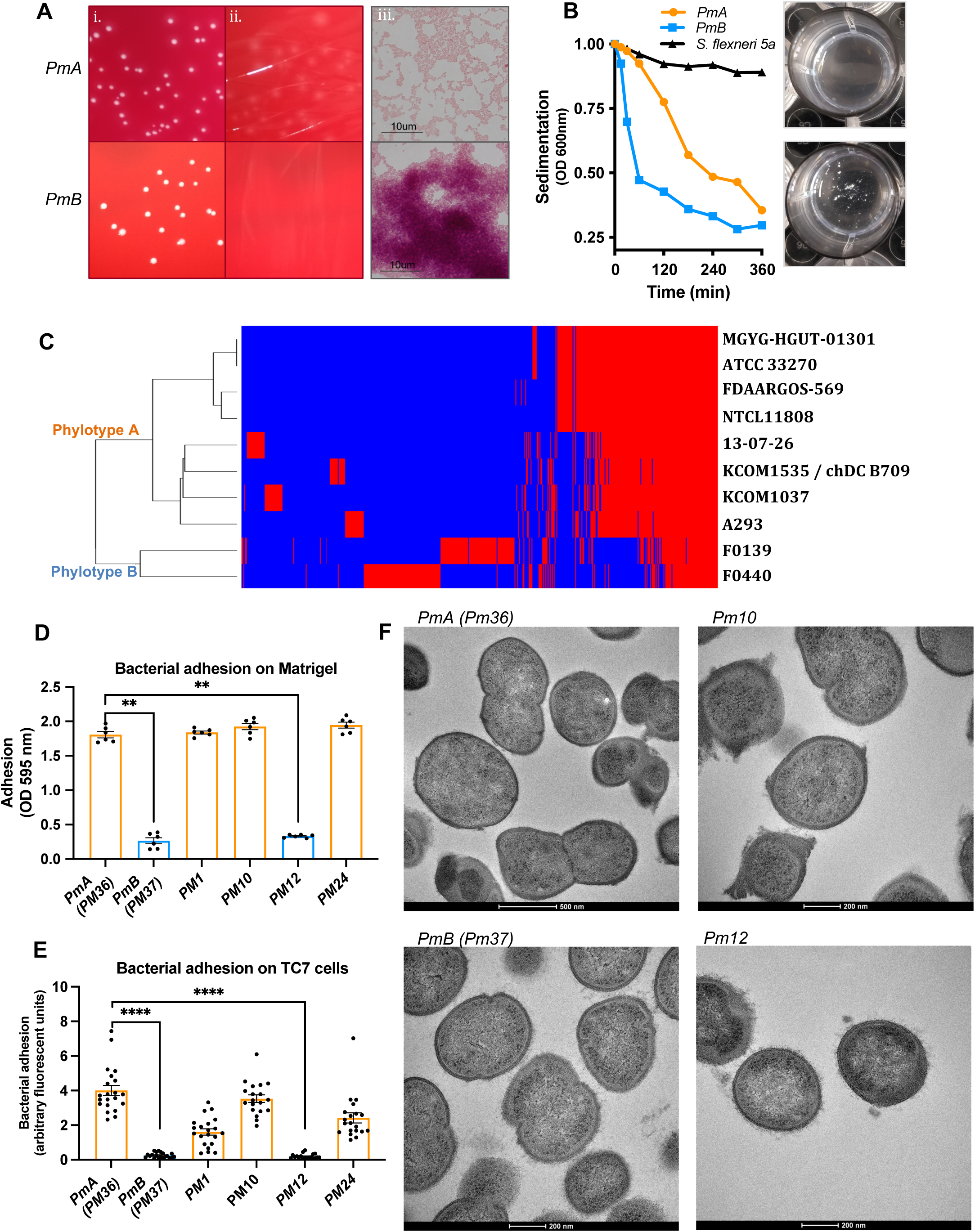
Phenotypic and genetic characterization of *P. micra*. A) Colony appearance of ATCC 33270 (*PmA*) (top) and HHM BlNa17 (*PmB*) (bottom) on blood agar plates: i) after 48 hours of culture under anaerobic conditions at 37°C; ii) when the colonies were removed to observe the hemolytic zone below the colonies; and iii) when colonies were analyzed via the Gram stain. B) Sedimentation assay performed on *PmA* and *PmB*. On the right, representative pictures of the suspension (top *PmA*, bottom *PmB*) obtained after 2 hours at 4°C without shaking. C) Clustering of *P. micra* phylotype A and B whole genomes according to the presence (red) or absence (blue) of genes. D) Adhesion capacity of *P. micra* ATCC 33270-*Pm36* (*PmA)* and HHM BlNa17-*Pm37* (*PmB)* and other oral isolates from phylotype A (*Pm1, Pm10 and Pm24*) or phylotype B (*Pm12*) to Matrigel®. Optical density at 595 nm represents measurement of bacterial adhesion. E) Adhesion capacity of *PmA*, *Pm1, Pm10, Pm24* from phylotype A and *PmB, Pm12* from phylotype B to human colonic cell lines TC7 after 1 hour of co-culture. Bacterial adhesion was quantified by the analysis of fluorescent images of cells co-cultured with the different isolates and is reported as arbitrary fluorescent units. Mann-Whitney test ****p<0.0001; **p<0.01. F). Transmission electron microscopy on ultrafine sections of phylotype A isolates *PmA* or *Pm10* (top), and phylotype B isolates *PmB* or *Pm12* (bottom).

**Figure S2:**
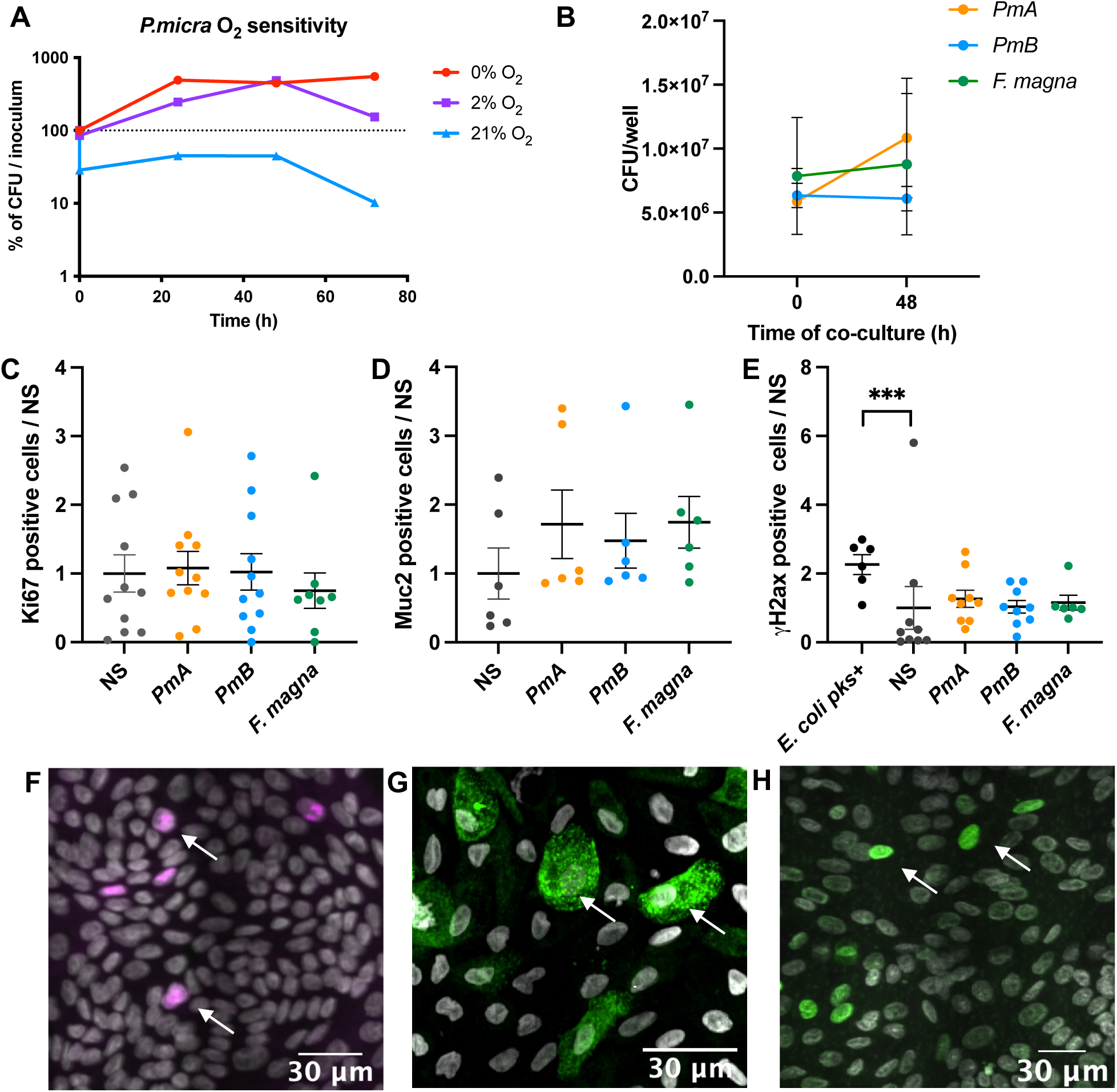
A) *P. micra* oxygen sensitivity. Representative growth curves of *P. micra* ATCC 33270 (*PmA*) under anaerobic (0% O_2_), aerobic (21% O_2_) and hypoxic (2% O_2_) conditions. At different incubation times, the cultures were plated on horse blood plates and returned to anaerobic conditions for 48 hours. CFUs were counted to estimate bacterial viability. The results are expressed as percentage of CFU obtained towards the inoculum grown at 0% O_2_. B) *P. micra* viability by CFU counts after 48 hours of co-culture with human colonic primary cells in hypoxic conditions (mean +/- SEM). C, D, and E. Quantification of proliferative cells, goblet cells, and cells harboring double-stranded DNA breaks by immunofluorescence using respectively an anti-Ki67, anti-Muc2, and anti-γH2ax labelling after 48 hours of co-culture with *P. micra* ATCC 33270 (*PmA*), *P. micra* HHM BlNa17 (*PmB*), *F. magna* ATCC 29328 or the *E. coli Pks*+ strain IHE3034 (used as a positive control for double-stranded breaks). NS, non-stimulated. Results are expressed as percentages of positive cells relative to the total number of nuclei and normalized using the non-treated sample (NS). Experiments were performed on cells from two donors, with at least 3 experiments per donor. The data are represented as mean +/- SEM. Mann-Whitney test ***p<0.001. F, G, and H are representative images of Ki67, Muc2 and γH2ax immunostaining, respectively.

**Table S1:**
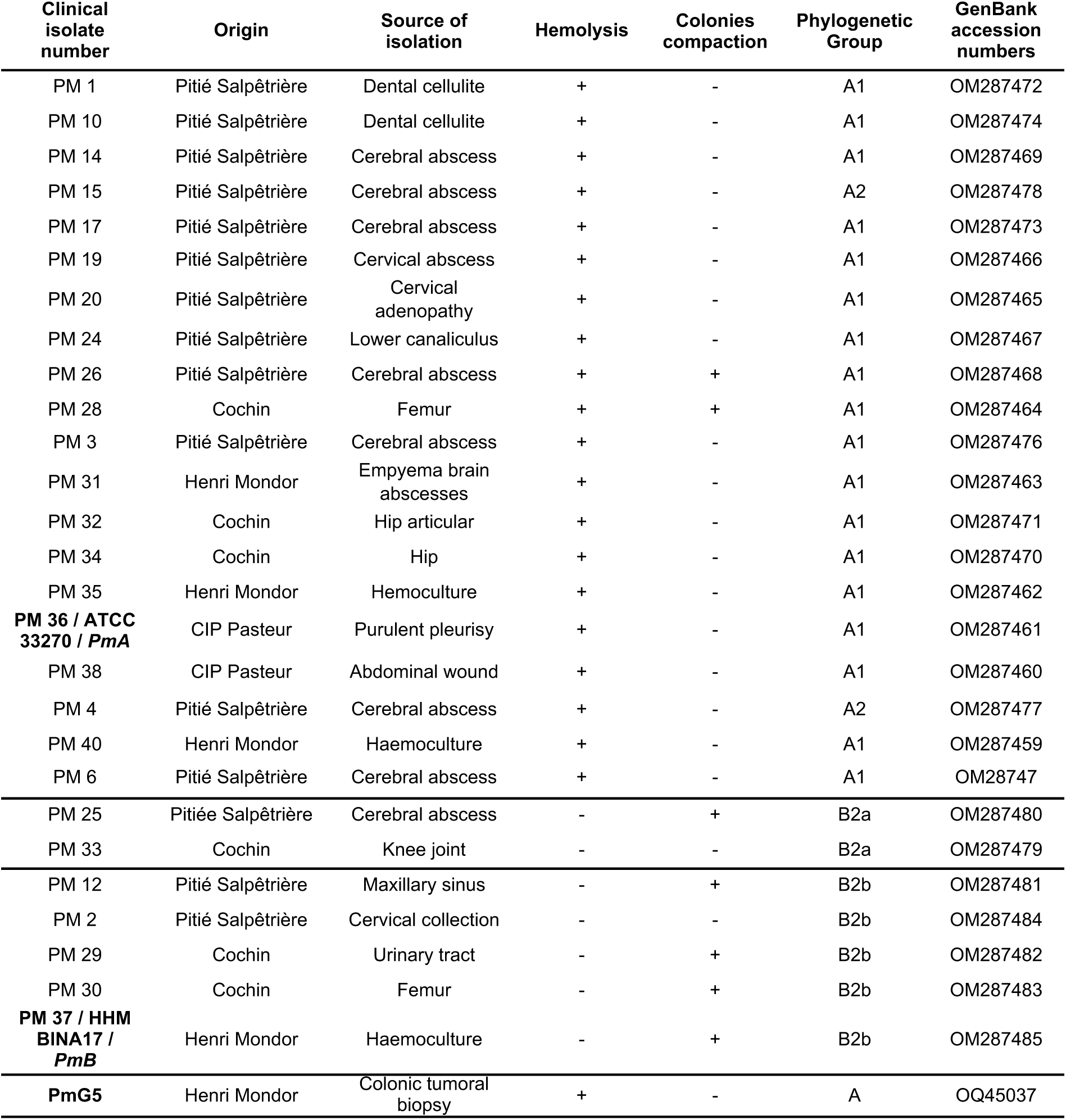
Description of *P. micra* clinical isolates obtained from several hospitals in Paris and from different infectious locations.

**Table S2:**
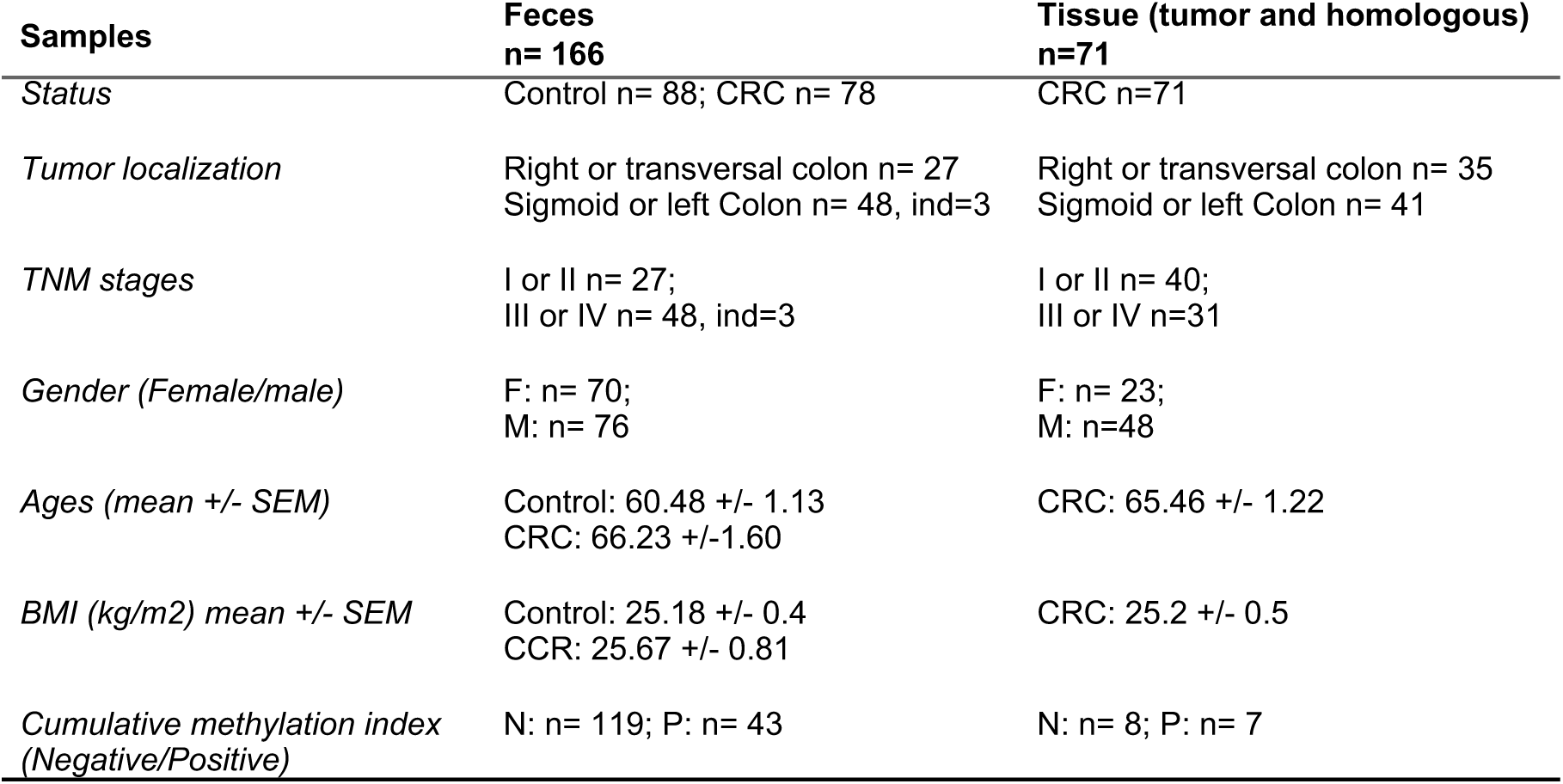
Clinical data.

**Table S3:** *PmA* induces DNA methylation changes in promoters of genes involved in carcinogenesis. CRC: colorectal cancer; TSG: tumor suppressor gene; EMT: epithelial-mesenchymal transition; IGFs: insulin-like growth factors; SCF: SKP1-CUL1-F-box protein; TRAPP: transport protein particle; ESCC: esophageal squamous cell carcinoma.

| Names, Alias | Described role | Link with CRC or cancers | References |
| --- | --- | --- | --- |
| **SCIN**<br>(Scinderin) | Ca(2+)-dependent actin-severing and capping protein. Regulation of actin cytoskeleton. | Overexpressed in CRC. Inhibits cell proliferation and tumorigenesis. | Lin 2019 |
| **HACE1**<br>(HECT domain E3 ubiquitin protein ligase 1) | E3 ubiquitin ligase involved in specific tagging of target proteins, leading to their subcellular localization or proteasomal degradation. | TSG. Down-regulated by DNA methylation in CRC. Loss or knockout of HACE1 enhanced tumor growth, invasion, and metastasis; in contrast, the overexpression of HACE1 can inhibit the development of tumors. | Li 2019<br>Hibi 2008<br>Zhang 2007 |
| **TSPAN13**<br>(Tetraspanin 13) | Transmembrane signal transduction protein that plays a role in the regulation of cell development, activation, growth, motility and invasion. | TSG. Downregulation inhibits proliferation of CRC cells. | Lou 2017 |
| **FBXO32**<br>(F-Box Protein 32)<br>Astagin-1 | Component of a SCF E3 ubiquitin-protein ligase complex which mediates the ubiquitination and subsequent proteasomal degradation of target proteins. | TSG. Under-expressed in CRC. Induce cell differentiation. Upstream regulator of EMT. | Yuan 2018<br>Sahy 2017 |
| **IGFBP7**<br>(Insulin Like Growth Factor Binding Protein 7) | Protein coding gene that regulate IGFs. Stimulates prostacyclin production and cell adhesion. Promotes cancer cell growth and migration. | TSG. Silencing induce metastasis. | Ruan 2007<br>Suzuki 2010 |
| **KIAA0494**<br>(EF-Hand Calcium Binding Domain 14) | Uncharacterized protein. Predicted membrane protein. | - |  |
| **DIAPH3**<br>(Diaphanous Related Fomrin 3) | Protein coding gene involve in actin remodeling. Regulate cell movement and adhesion. | Deficiency enhances cell motility, invasion and metastasis in many cancers. | Hager 2012<br>Rana 2018 |
| **SIX1**<br>(sine oculis homeobox 1) | Transcription factor involve in regulation of cell proliferation, apoptosis, embryonic development and tumorigenesis. | Oncogene. Overexpressed in CRC, overexpression of Six1 dramatically promotes CRC tumor growth and metastasis in vivo. | Wu 2014<br>Xu 2017 |
| **KCNB1**<br>(Potassium Voltage-Gated Channel Subfamily B Member 1) | Voltage-gated potassium channel Kv2.1. Contributes to the pronounced pro-apoptotic potassium current surge during neuronal apoptotic cell death in response to oxidative injury. | KCNB1 polymorphisms correlate to CRC treatment and patient's outcome. Kv2.1 occasionally forms complexes with other voltage-gated potassium alpha-subunits (i.e. Kv9.3), witch its silencing potently inhibiting proliferation in human colon cells. | Li 2015,<br>Fahra 2020 |
| **SASH1**<br>(SAM and SH3 domain-containing protein 1) | Scaffold protein involved in the TLR4 signaling. Stimulate cytokine production and endothelial cell migration in response to binding pathogens | TSG. Downregulated in CRC. Downregulation expression was correlated with the formation of metachronous distant metastasis. Loss of SASH1 induces EMT. SASH1 inhibits metastasis formation In Vivo. | Rimkus 2006,<br>Franke 2019 |
| **ZNF282**<br>(Zinc Finger Protein 282) | Transcription factor binding the U5RE (U5 repressive element) of HLTV-1 (human T cell leukemia virus type 1) with a repressive effect. Co-activator of the estrogen receptor alpha. | Knockdown reduced migration, invasion and tumorigenesis of ESCC *in vitro* and reduced the tumorigenicity of ESCC xenograft in nude mouse. | Yeo 2014 |
| **SEMA3F**<br>(Semaphorin 3F) | Secreted signaling protein that are involved in axon guidance during neuronal development. Act in an autocrine fashion to induce apoptosis, inhibit cell proliferation and survival. Regulator of actin cytoskeleton. | TSG. Down-regulated by DNA methylation in CRC tissues and CRC cell lines. Associated with progressive phenotypes of CRC. Overexpression reduced proliferation, adhesion, and migratory capability of colon cancer cells. SEMA3F-overexpressing cells exhibited diminished tumorigenesis when transplanted in nude mice and reduced liver metastases. | Gao 2015,<br>Wu 2011,<br>Bielenberg 2004 |

**Tables S4:** Excel file with i) the list of differentially expressed genes (DEGs) between 1- *Parvimonas and 2- P.micra phylotype A*–negative and – positive patients; ii) Pathway enrichment analysis on RNaseq results; iii) List of differentially methylated regions (DMRs) between *P. micra* –negative and – positive patients.

